# UPR-induced intracellular C5aR1 promotes adaptation to the hypoxic tumour microenvironment by regulating tumour cell fate

**DOI:** 10.1101/2024.09.27.615431

**Authors:** Tatsuya Suwa, Kelly Lee, Ian Chai, Heather Clark, David J. MacLean, Nicole Machado, Gonzalo Rodriguez Berriguete, Geoff S Higgins, Ester M Hammond, Monica M Olcina

## Abstract

Dysregulation of the C5a-C5a receptor 1 (C5aR1) signalling axis underlies inflammation and immune-driven pathology. C5aR1 was traditionally thought to be primarily expressed on the cell membrane, although recent reports indicate the importance of intracellular C5aR1 expression for the inflammatory effector functions of various cell types. However, the mechanisms regulating C5aR1 expression and localisation remain unclear. In tumours with an immunosuppressive microenvironment, we recently found robust C5aR1 expression on malignant epithelial cells, highlighting potential tumour cell–specific functions. Here, we show that hypoxia, a hallmark of immunosuppressive microenvironments, induces C5aR1 expression in an unfolded protein response (UPR)-dependent manner via enhanced endoplasmic reticulum stress. Furthermore, hypoxia drives endocytosis, relocating C5aR1 from the cell membrane to the intracellular compartment. By genetically and pharmacologically targeting the C5a/C5aR1 axis, we show that C5aR1 mediates cellular adaptation to hypoxia by regulating processes associated with cell fate, including autophagy and apoptosis. In line with hypoxia-induced intracellular C5aR1 pools, pharmacological inhibition of C5aR1, particularly with cell-permeable small molecule inhibitors, significantly reduces tumour cell survival. These results suggest that the dysregulated C5a/C5aR1 axis and the hypoxia-induced shift in C5aR1 localisation, support tumour cell survival and provide new insights into therapeutic strategies for targeting the C5a/C5aR1 axis.

## Introduction

A range of low oxygen concentrations (hypoxia) are a key feature of the tumour microenvironment (TME). Hypoxia is caused by heterogeneity of tumour oxygenation resulting from weak vascular structures and constantly proliferating cells. Accumulating evidence has shown that tumour hypoxia, is associated with poor prognosis in patients with solid tumours ^1–3^.

The biological response to hypoxia is well-known to result in immunosuppression and treatment resistance, which contributes to the poor outcome of tumours with hypoxic TMEs ^3–5^. We recently reported that complement component C5a receptor 1 (C5aR1) can cause treatment resistance following radio- and chemotherapy in tumours with an immunosuppressive TME ^6^. The complement system is an ancient innate immune pathway with systemic and local immune functions. Previous studies have reported that hypoxia contributes to complement dysregulation in the TME via HIF-dependent upregulation of negative regulators of the complement system which are highly expressed in hypoxic cancer cells and may limit complement-dependent cytotoxicity ^7–9^.

Enhanced local production of complement anaphylatoxins such as the C5aR1 ligand C5a, by various cells in the TME, following chronic inflammation or via treatment-induced cell death, can also contribute to complement dysregulation in cancer. Moreover, recent studies on non-canonical functions for complement components have reported cell autonomous expression of complement components and receptors in several cancer cells. Cell autonomous signalling via complement receptors has been shown to activate intracellular signalling pathways such as WNT/β-catenin, NFκB, and PI3K/AKT, leading to malignant characteristics including proliferation, invasion and metastasis ^10–14^. In addition, it has been recently reported that intracellular cleavage of C3 and C5 occurs in cancer cells as well as immune cells such as T cells and macrophages ^12,15,16^. However, the mechanism regulating expression and cellular localisation of complement receptors such as C5aR1 in the TME remains unclear.

Under levels of hypoxia associated with treatment resistance and immunosuppression (<0.1% O_2_), protein folding in the endoplasmic reticulum (ER) is inhibited. This leads to an accumulation of misfolded and unfolded proteins in the ER lumen, a condition called ‘ER stress’ ^17^. The unfolded protein response (UPR) plays an important role in the cytoprotective adaptation that reduces ER stress and maintains protein homeostasis ^18,19^. The UPR consists of three signalling pathways initiated by the following three ER-localised signal sensors; protein kinase R–like endoplasmic reticulum kinase (PERK), inositol requiring enzyme 1 (IRE1) and activating transcription factor 6 (ATF6). In unstressed conditions, 78-kDa glucose-regulated protein (GRP78), an ER chaperone also termed binding immunoglobulin protein (BiP), binds to these signal sensors and prevents the activation of their downstream signalling, thereby maintaining the UPR signalling in an inactive state. In hypoxic conditions (<0.1% O_2_), the UPR signalling becomes activated as BiP dissociates from these sensors to activate their downstream signalling cascade. Accumulating evidence has shown that activation of UPR signalling regulates autophagy and apoptosis ^18,20^, as well as the degradation of misfolded proteins, the biosynthesis of amino acids and lipids, and redox homeostasis, to reduce cellular stress and restore cellular homeostasis ^21,22^. These UPR functions mediate the adaptation of cancer cells to ER stress under hypoxia, leading to tumour progression ^23,24^. Here, we show that cancer cells under hypoxia and ER stress increase the expression of intracellular C5aR1 in a UPR-dependent manner. Increased C5aR1 expression results in attenuated autophagy and apoptosis thereby increasing cancer cell survival under hypoxia (<0.1% O_2_). Furthermore, we also show that genetic and pharmacological inhibition of C5aR1 signalling decrease cancer cell survival under hypoxic conditions, with strategies able to inhibit intracellular C5aR1 pools showing the greatest effects on survival. Collectively, our findings provide new insights into the regulatory mechanisms and functions of C5aR1 and highlight how this information can be used to therapeutically target the most therapy-resistant populations of the TME.

## Results

### C5aR1 is highly expressed in hypoxic regions and is associated with poor outcome in various cancers

Our work and that of others has previously shown that the expression of C5aR1 is higher in tumours compared to normal tissues in several cancers such as gastric, colorectal, and cervical cancer ^25–27^. We investigated C5aR1 expression in tumour and normal tissues with an in-house tissue microarray (TMA) consisting of tumour and normal tissue pairs. Consistent with previous studies, we found that, at the protein level, C5aR1 was highly expressed in tumour compared to normal tissues in all of prostate and endometrium sections, and three out of five colorectal sections (Figure 1A). We next analysed the significance of C5aR1 expression on prognosis in various types of cancers using the TCGA database and found that high C5aR1 mRNA expression is significantly associated with poor prognosis in various cancers, including colon, prostate, ovarian cancer, and glioblastoma (Figure 1B and Supplementary figure S1A).

**Figure 1.**
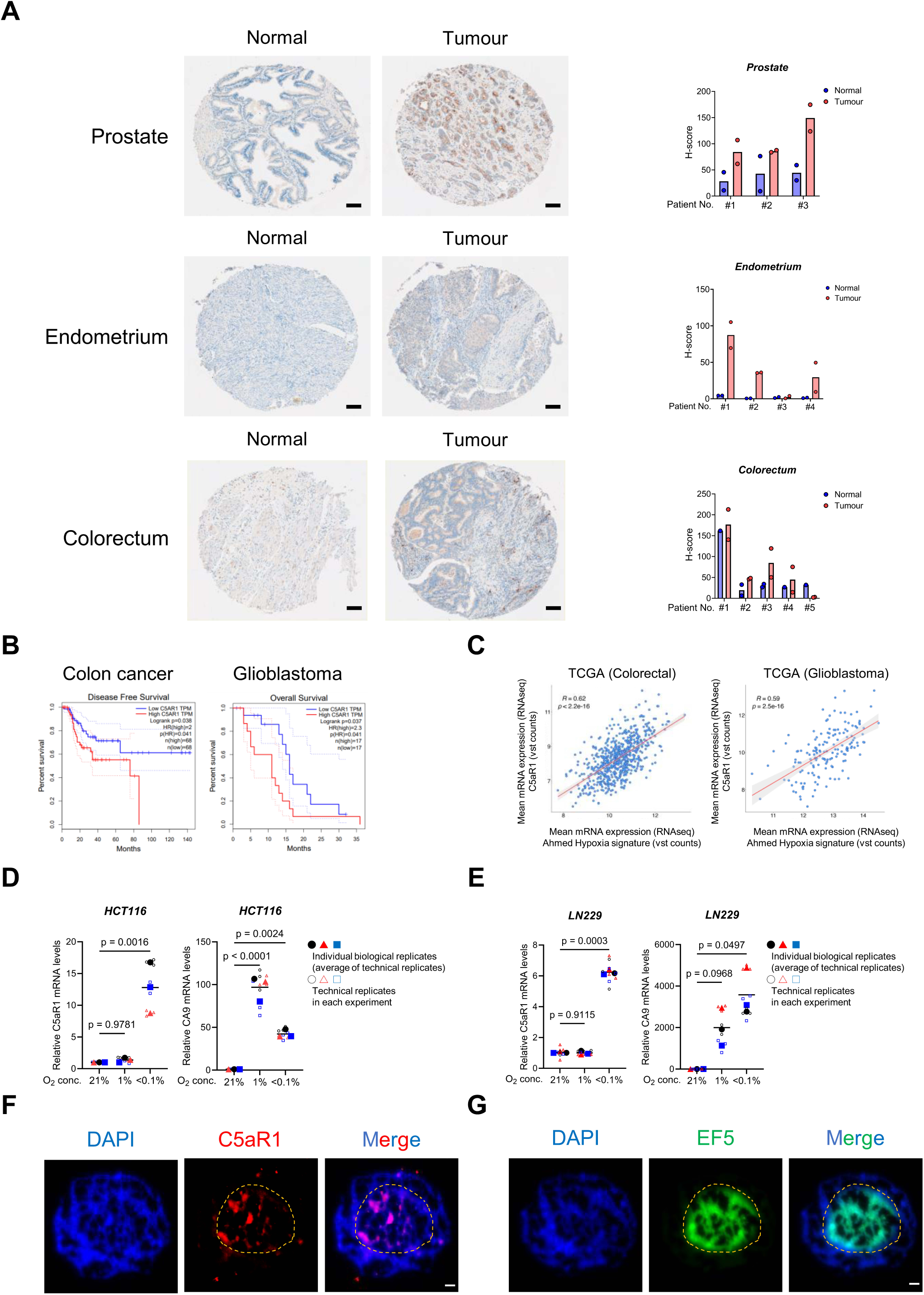
C5aR1 is highly expressed in hypoxic regions and is associated with poor outcome in various cancers. **(A)** Representative images of C5aR1 staining in TMA for prostate, endometrial, and colorectal normal and tumour tissues. Scale bar, 100 µm. H-score is shown. 1 or 2 cores per tissue.**(B)** GEPIA Kaplan-Meier curve for disease free survival of TCGA colon cancer patients (left) and overall survival of TCGA glioblastoma patients (right) with high (red) or low (blue) C5aR1 mRNA expression levels is shown. (http://gepia.cancer-pku.cn). **(C)** Pearson’s correlation of C5AR1 mRNA expression and Ahmed Hypoxia signature in TCGA colorectal cancer and glioblastoma samples. *R* score and *p* value are shown. **(D and E)** HCT116 (D) and LN229 (E) cells were cultured under the indicated oxygen conditions for 24 hours (hr) and 16 hr, respectively, and subjected to qRT-PCR. n=3. Individual biological replicates (large points) represent the average of the technical replicates (small points). *p* values were calculated using biological replicates by one-way ANOVA with Dunnett’s test. **(F and G)** Serial sections of HCT116 spheroids treated with EF5 were stained with the indicated antibodies; C5aR1 (red), a hypoxia marker, EF5 (green), or DAPI (blue). The dotted line represents the estimated outside edge of the EF5-positive regions. Scale bar, 50 µm.

We recently reported that in tumour models with immunosuppressive TMEs, C5aR1 is highly expressed in malignant epithelial cells ^6^. We hypothesised that the increased expression of C5aR1 may contribute to adaptation to stress within the TME. As tumour hypoxia is a key feature of the TME in solid tumours and is strongly associated with immunosuppression ^3,28^, we investigated the possibility that the dysregulation of C5aR1 may be caused by hypoxia-induced stress. TCGA analysis indicated that C5aR1 mRNA levels were significantly correlated with hypoxia signatures in glioblastoma, colorectal and prostate cancer, suggesting that C5aR1 expression may be increased in the hypoxic environment of tumours (Figure 1C and Supplementary figure S1B). To investigate the effect of hypoxia on C5aR1 expression, we examined C5aR1 mRNA levels at different oxygen concentrations. C5aR1 mRNA levels were significantly increased in severe hypoxia (<0.1% O_2_) but not milder levels (1% O_2_) in colorectal and glioblastoma cancer cells (Figure 1D and 1E, and Supplementary figure S1C-S1F, showing similar effect in ovarian and prostate cancer cells). qPCR results showed that hypoxia also increases mRNA levels of C5 (which gives rise to C5aR1’s ligand C5a) in an oxygen-dependent, with increased expression observed only under severe hypoxia in HCT116 colorectal cancer cells (Supplementary figure S1G and S1H). To investigate the relationship between hypoxia and C5aR1 expression in a 3D model, we prepared spheroids treated with EF5, a hypoxia marker ^29^, and performed immunohistochemical analyses. C5aR1 was expressed in the EF5-positive regions of the spheroids, suggesting again that C5aR1 was induced by hypoxia (Figure 1F and 1G). Serial sections (rather than co-localisation in a single section) are shown here since both the C5aR1 and anti-EF5 antibody are raised in the same species. Taken together, these results indicate that hypoxia (<0.1% O_2_) upregulates expression of C5aR1.

### ER stress induces C5aR1 expression in cancer cells under hypoxia (<0.1% O_2_)

Next, we investigated the molecular mechanisms underlying the upregulation of C5aR1 expression under hypoxia (<0.1% O_2_). We first examined the influence of HIF-1α, the master transcription factor that regulates gene expression under hypoxia ^30,31^. *In vitro* experiments showed that the increase in C5aR1 mRNA levels under hypoxia was still observed in HIF-1α-knock out (KO) and WT cells suggesting that hypoxia induces C5aR1 mRNA expression in a HIF-1α-independent manner (Figure 2A, and Supplementary figure S2A). In addition, treatment with HIF-inducer cobalt chloride (CoCl_2_) did not result in increased C5aR1 mRNA levels (Figure 2B, and Supplementary figure S2B). These data are consistent with our data showing that C5aR1 is not significantly induced under milder hypoxic conditions (1% O_2_) associated with a robust HIF-mediated response. Since p53 is a driver of gene expression regulation under severe hypoxia (<0.1% O_2_) ^32^, we assessed the effect of p53 in regulating hypoxia-induced C5aR1 expression. qPCR results showed that the increase in C5aR1 mRNA levels under hypoxia was still observed in p53 KO cells (Supplementary figure S2C). These results suggest that the hypoxia-dependent increase in C5aR1 mRNA levels is independent of HIF-1α and p53.

**Figure 2.**
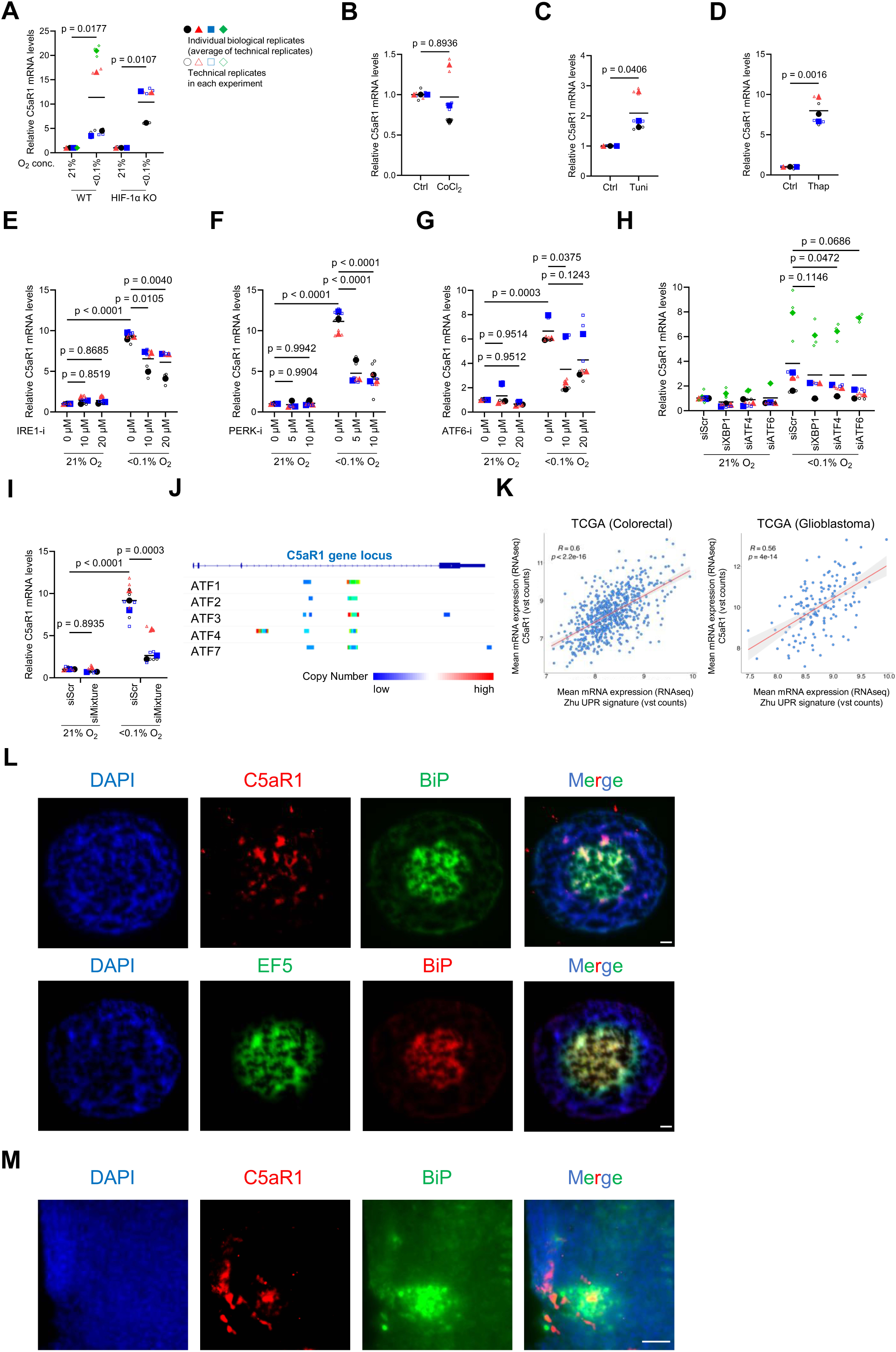
ER stress induces C5aR1 expression in cancer cells under hypoxia (<0.1% O_2_) For the whole figure: Individual biological replicates (large points) represent the average of the technical replicates (small points). *p* values were calculated using biological replicates by two-tailed paired Student’s t test (A), two-tailed unpaired Student’s t test (B-D), two-way ANOVA with uncorrected Fisher’s LSD test (E-G and I), or one-way ANOVA with Dunnett’s test (H). **(A)** HIF-1α-KO and WT HCT116 cells were cultured under normoxia or hypoxia (<0.1% O_2_) for 24 hr and subjected to qRT-PCR. n=3 and 4. **(B-D)** HCT116 cells were treated with either 200 µM cobalt chloride (CoCl_2_) for 24 hr (B), 10 µg/mL tunicamycin (Tuni) for 24 hr (C), 2 µM thapsigargin (Thap) for 16 hr (D), or its vehicle (Ctrl), and subjected to qPCR. n=3. **(E-G)** HCT116 cells were cultured under normoxia or hypoxia (<0.1% O_2_) in the presence of either IRE1-inhibitor (E), PERK-inhibitor (F), or ATF6-inhibitor (G), and subjected to qPCR. n=3. **(H)** HCT116 cells were transfected with the indicated siRNA or scramble siRNA (siScr), cultured under normoxia or hypoxia (<0.1% O_2_) for 24 hr, and subjected to qPCR. n=4. **(I)** After simultaneously silencing XBP1, ATF4, and ATF6 using siRNA mixtures, HCT116 cells were cultured under normoxia or hypoxia (<0.1% O_2_) for 24 hr, and subjected to qPCR. n=3. **(J)** Enrichment Analysis with public Chip-seq data on Chip-Atlas website is shown (https://chip-atlas.org/peak_browser). Colours represent copy numbers of the indicated transcription factors binding to C5aR1 gene locus in all cell lines. **(K)** Pearson’s correlation of C5AR1 mRNA expression and Xhu UPR signature in TCGA colorectal cancer and glioblastoma samples. *R* score and *p* value are shown. **(L and M)** Serial sections of HCT116 spheroids treated with EF5 (L) or HCT116 tumour xenografts (M) were stained with the indicated antibodies; (L) Section 1, C5aR1(red), BiP (green), or DAPI (blue); Section 2, BiP (red), EF5 (green), or DAPI (blue). (M) C5aR1 (red), BiP (green), or DAPI (blue). Scale bar, 50 µm.

In line with previous reports^17,22,33^, we observed that the UPR-related gene, CHOP, was significantly up-regulated specifically under severe hypoxia (Supplementary figure S2D), supporting the evidence that the UPR is a cellular ER stress response mounted under severe hypoxia. We hypothesised that ER stress may induce C5aR1 expression, and found that C5aR1 mRNA levels were increased in the presence of two different UPR inducers, tunicamycin and thapsigargin (Figure 2C and 2D, and Supplementary figure S2E-H). Interestingly, C5 mRNA levels were also increased following treatment with UPR inducers in colorectal HCT116 cells (Supplementary figure S2I and S2J). In the context of UPR signalling, IRE1, PERK and ATF6 are three key UPR signal activator proteins ^17,18^. We, therefore, next examined whether these UPR activator proteins were responsible for the induction of C5aR1 expression under hypoxia. Interestingly, qPCR results showed that the induction of C5aR1 mRNA levels under hypoxia was partially but significantly suppressed in the presence of UPR inhibitors against IRE1, PERK, and ATF6 (Figure 2E-2G, and Supplementary figure S2K-S2M). These results suggest that in hypoxia each of these pathways is involved in the induction of C5aR1 expression. To further narrow down the involvement of these pathways, we examined the effects of XBP1, ATF4 and ATF6, the major transcription factors downstream of IRE1, PERK and ATF6, respectively. Silencing of either XBP1, ATF4 or ATF6 alone showed a trend towards suppression, but not complete inhibition of C5aR1 mRNA induction under hypoxia, suggesting that the UPR pathways act in a redundant manner to regulate C5aR1 (Figure 2H and Supplementary figure S2N). To test this possibility, we examined the effect of simultaneous silencing of the three representative UPR pathways on the induction of C5aR1 mRNA under hypoxia. The induction of C5aR1 mRNA was decreased by simultaneous silencing of XBP1, ATF4 and ATF6, suggesting that these three pathways act complementarily to induce C5aR1 expression under hypoxia (Figure 2I and Supplementary figure S2O). Furthermore, querying publicly available ChIP datasets indicated that several UPR-related transcription factors including ATF4 bind to the C5aR1 gene locus, again suggesting that the UPR regulates C5aR1 expression via multiple transcription factors (Figure 2J).

We next assessed the relationship between C5aR1 expression and UPR signalling in clinical samples. TCGA analysis showed that C5aR1 mRNA levels are strongly correlated with a UPR signature in various types of tumours including glioblastoma, colorectal, ovarian and prostate cancers (Figure 2K and Supplementary figure S2P). Further analysis indicated that the hypoxia signature used in figure 1C strongly correlated with this UPR signature (Supplementary figure S2Q). These data suggest that C5aR1 is expressed in regions of the tumour associated with the biological response to hypoxia and UPR stress. As we observed that C5aR1 is expressed in EF5-positive regions in the HCT116 colorectal cancer spheroids (Figure 1F and 1G), we then examined whether C5aR1 was expressed in BiP (an intrinsic UPR marker)-expressing areas. As expected, C5aR1 positive areas were associated with BiP-expressing areas in HCT116 spheroids (Figure 2L). In addition, we also observed that C5aR1 was expressed in BiP-expressing areas in HCT116 xenografts (Figure 2M). Taken together, these results suggest that expression of C5aR1 is induced in a UPR-dependent manner in response to ER stress under severe hypoxia.

### Hypoxia-induced C5aR1 mediates cellular adaptation to hypoxic stress by regulating cancer cell death

During cellular stress, the UPR pathway is known to impact cell fate and adaptation to stress by regulating autophagy and apoptosis ^17,18,20^. We, therefore, asked whether hypoxia-induced-UPR might mediate adaptation to ER stress and return to homeostasis via C5aR1-regulated autophagy and/or apoptosis. We found that under hypoxic conditions, autophagic flux as assessed by decreased levels of p62, a selective autophagy receptor which is degraded when autophagy is induced, was increased by C5aR1 silencing (Figure 3A, and Supplementary figure S3A and S3B), and decreased by C5aR1 overexpression (Figure 3B, and Supplementary figure S3C and S3D). In addition, apoptotic cell death was increased under hypoxia by silencing of C5aR1 (Figure 3C-D, and Supplementary figure S3E and S3F). To further assess the impact of hypoxia-mediated dysregulation of the C5a-C5aR1 axis, we next examined the influence of silencing C5 and noted similar changes in autophagy (Figure 3E, and Supplementary figure S3G) and apoptosis (Figure 3F, and Supplementary figure S3H) as following genetic manipulation of C5aR1. These results suggested that a dysregulated C5-C5aR1 axis impacts autophagy and apoptosis under hypoxic stress.

**Figure 3.**
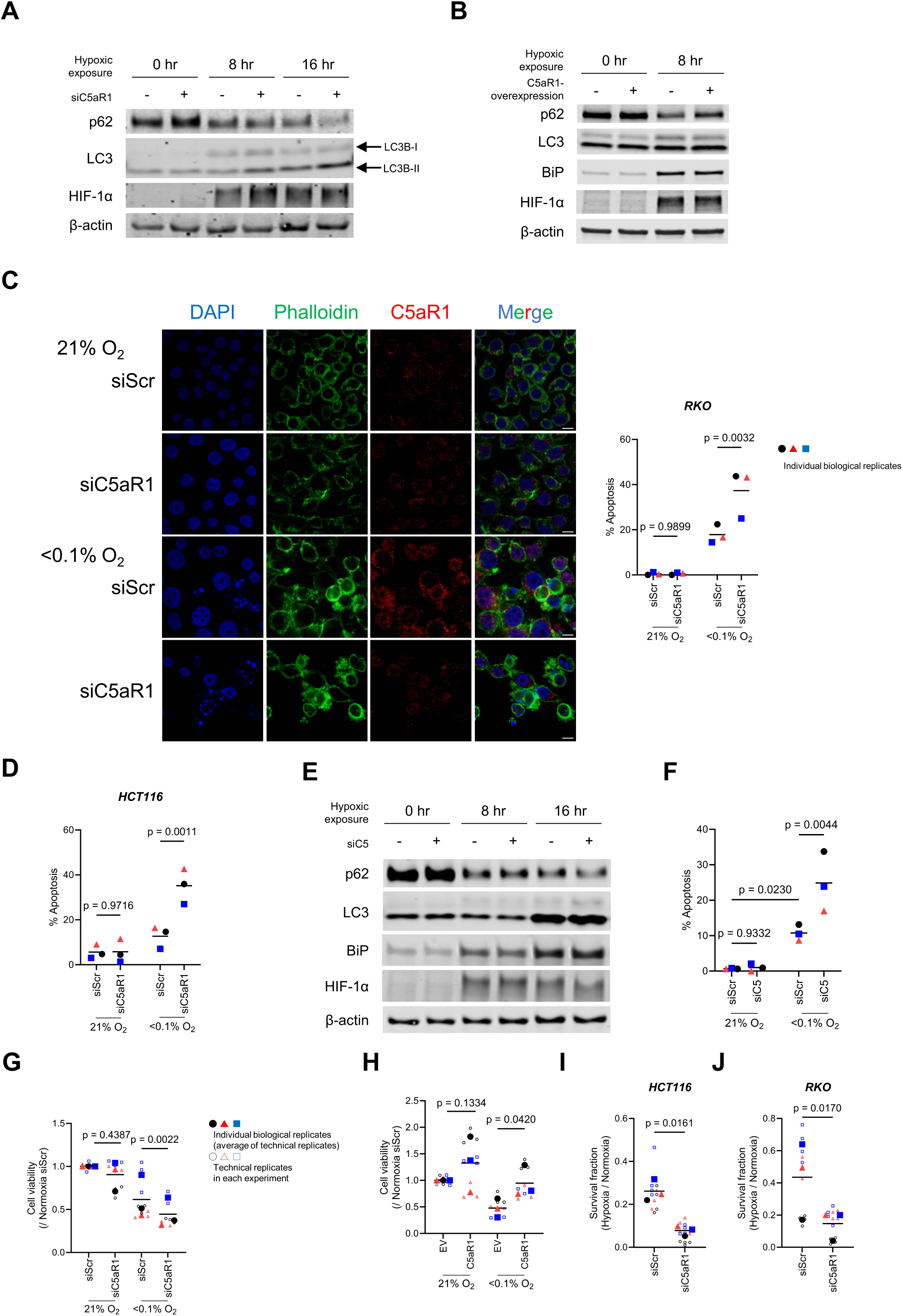
Hypoxia-induced C5aR1 mediates cellular adaptation to hypoxic stress by regulating cancer cell death. For the whole figure: Individual biological replicates (large points) represent the average of the technical replicates (small points)*. p* values were calculated using biological replicates (large points) by two-way ANOVA with uncorrected Fisher’s LSD test (C, D, and F) or two-tailed paired Student’s t test (G-J). **(A and B)** HCT116 cells were transfected with either siC5aR1 or siScr (A) or with either pcDNA3.1/C5aR1-GFP (C5aR1) or its empty vector (EV) (B), cultured under normoxia or hypoxia (<0.1% O_2_) for the indicated periods, and subjected to Immunoblotting. **(C)** RKO cells were transfected with siC5aR1 or siScr, cultured under normoxia or hypoxia (<0.1% O_2_) for 24 hr, and subjected to immunocytochemistry and apoptosis assay. C5aR1 (red), Phalloidin (green), or DAPI (blue). Scale bar, 10 µm. n=3. **(D)** HCT116 cells were transfected with either siC5aR1 or siScr, cultured under normoxia or hypoxia (<0.1% O_2_) for 24 hr and subjected to apoptosis assay. n=3. **(E)** HCT116 cells were transfected with either siRNA against C5 (siC5) or siScr, cultured under normoxia or hypoxia (<0.1% O_2_) for the indicated periods, and subjected to immunoblotting. **(F)** RKO cells were transfected with either siC5 or siScr, cultured under normoxia or hypoxia (<0.1% O_2_) for 24 hr, and subjected to apoptosis assay. n=3. **(G and H)** HCT116 cells were transfected with either siC5aR1 or siScr (G) or with either pcDNA3.1/C5aR1-GFP (C5aR1) or EV (H), cultured under normoxia or hypoxia (<0.1% O_2_) for 40 hr (G) and 32 hr (H), respectively, and subjected to cell viability assay. n=3. **(I and J)** HCT116 (I) and RKO (J) cells were transfected with either siC5aR1 or siScr, cultured under normoxia or hypoxia (<0.1% O_2_) for 24 hr and subjected to clonogenic survival assay. n=3.

The UPR pathway is also involved in the DNA damage and antioxidant responses, as well as the regulation of DNA replication stress, and cell cycle ^22,34–36^. However, silencing of C5aR1 did not affect cell cycle distribution, ROS production or DNA damage and replication stress markers (Supplementary figure S3I-K).

Next, we directly assessed the role of hypoxia-induced C5aR1 on cell survival. We found that, under severe hypoxia, cell viability was significantly reduced by C5aR1 knockdown (Figure 3G), whereas it was significantly increased by C5aR1 overexpression (Figure 3H). Consistent with the cell viability assay, the clonogenic survival assay showed that silencing of C5aR1 significantly decreased cell survival under hypoxia (Figure 3I and 3J, and Supplementary figure S3L). Collectively, these results indicate that C5aR1 mediates cellular adaptation to hypoxic stress by modulating autophagy and apoptosis in cancer cells.

### Pharmacologically targeting intracellular C5aR1 results in reduced tumour cell viability with enhanced autophagy and apoptosis

Having shown that genetically suppressing hypoxia-induced C5aR1 expression decreased cell viability and cell survival, we then investigated the impact of pharmacological targeting the C5a-C5aR1 axis. We first tested PMX205, a selective C5aR1 antagonist extensively used preclinically and currently undergoing clinical testing for amyotrophic lateral sclerosis (ALS) ^37,38^. *In vitro*, the effect of PMX205 resulted in a significant but very modest reduction in cell survival in hypoxia (Figure 4A and Supplementary figure S4A). This is in line with our previous data showing that PMX205 does not significantly delay tumour growth in colorectal tumour models *in vivo* unless used in combination with other cytotoxic therapies ^6^. We next used other selective C5aR1 inhibitors, JPE-1375 and Avacopan ^39,40^ and found that both of these showed more significant effects on cancer cell viability than PMX205 (Figure 4B and Supplementary figure S4B). We examined whether the effects on cell survival were associated with enhanced autophagy and apoptosis and found that, consistent with cell viability, JPE-1375 and Avacopan showed greater effects than PMX205 (Figure 4C and 4D).

**Figure 4.**
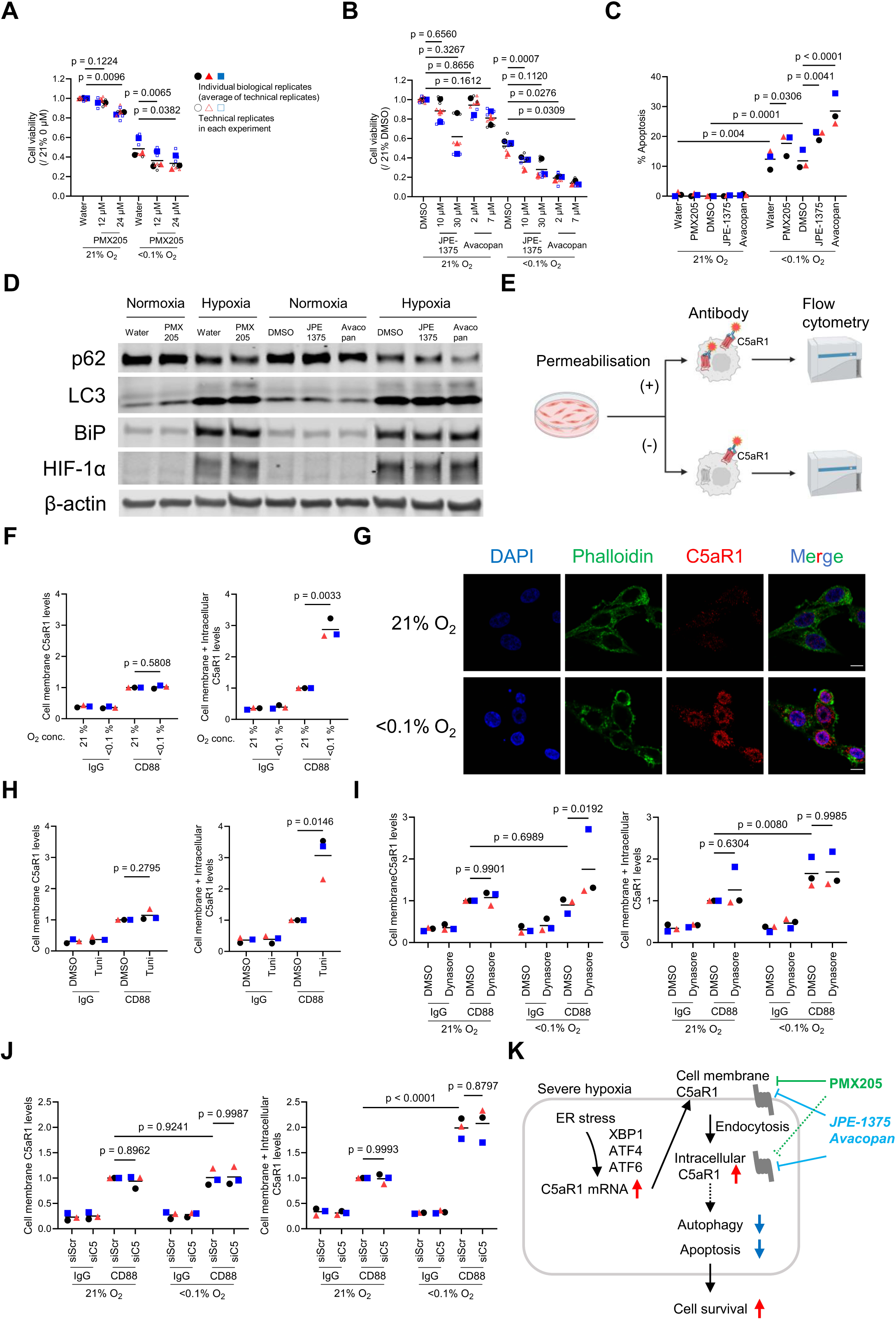
Pharmacologically targeting intracellular C5aR1 results in reduced tumour cell viability with enhanced autophagy and apoptosis. For the whole figure: Individual biological replicates (large points) represent the average of the technical replicates (small points)*. p* values were calculated using biological replicates (large points) by one-way ANOVA with Dunnett’s test (A and B), two-way ANOVA with uncorrected Fisher’s LSD test (C, I and J), and two-tailed paired Student’s t test (F and H). **(A-B)** HCT116 cells were pretreated for 8 hr with the indicated dose of C5aR1 antagonists, PMX205, JPE-1375 and Avacopan. Cells were cultured under normoxia or hypoxia (<0.1% O_2_) for 40 hr and subjected to cell viability assays. n=3. **(C-D)** HCT116 cells were pretreated for 8 hr with 12 μM PMX205, 10 μM JPE-1375 and 2 μM Avacopan. Cells were cultured under normoxia or hypoxia (<0.1% O_2_) for 24 hr and 16 hr and were subjected to apoptosis assay (C) and immunoblotting (D), respectively. n=3. (**E**) Schematic representation of experimental design for F and H–J. Created in BioRender.com. (**F**) RKO cells were cultured under normoxia or hypoxia (<0.1% O_2_) for 16 hr, and subjected to FACS, with (right) or without (left) permeabilisation. n=3. (**G**) RKO cells were cultured under normoxia or hypoxia (<0.1% O_2_) for 24 hr, and subjected to triple immunofluorescence. C5aR1 (red), Phalloidin (green), or DAPI (blue). Scale bar, 10 µm. (**H**) After treatment with 10 µg/mL tunicamycin (Tuni) or vehicle (DMSO) for 24 hr, HCT116 cells were subjected to FACS with (right) or without (left) permeabilisation. n=3. (**I**) HCT116 cells were cultured under normoxia or hypoxia (<0.1% O_2_) in the presence of 100 µM Dynasore and subjected to FACS with (right) or without (left) permeabilisation. n=3. (**J**) HCT116 cells were transfected with either siRNA against C5 (siC5) or siScr, cultured under normoxia or hypoxia (<0.1% O_2_) for 24 hr, and subjected to FACS with (right) or without (left) permeabilisation. n=3. (**K**) Working model: In cancer cells, UPR-induced C5aR1 is internalised and accumulated by endocytosis under hypoxia. Intracellular C5aR1 contributes to cancer cell survival by modulating autophagy and apoptosis under hypoxia. To effectively target the C5a/C5aR1 axis in the TME, cell permeable C5aR1 inhibitors may be more effective.

As a G-protein coupled receptor (GPCR), C5aR1 is typically thought to be expressed on the cell membrane ^8,10,41,42^. However, recent studies have reported that C5aR1 is also expressed intracellularly ^12,15,43^. Interestingly, unlike PMX205, JPE-1375 and Avacopan are predicted to be cell permeable ^37,39,40,43–45^. Given that these agents had no significant effect under normoxia but led to increased cell death under hypoxia (Figure 4A and 4B, and Supplementary figure S4A and S4B), we hypothesised that hypoxia may alter not only the expression but also the subcellular localisation of C5aR1, and that this might impact the efficacy of different pharmacological approaches to target C5aR1. Since there is no specific C5aR1 antibody available for immunoblotting, we initially investigated this hypothesis using flow cytometry (with or without permeabilisation) (Figure 4E). These studies indicated that under hypoxic conditions there is an increase in total C5aR1, but not cell membrane C5aR1 expression, suggesting that it is the intracellular C5aR1 pool which increases under hypoxia (Figure 4F, and Supplementary figure S4C and S4D). We then performed immunofluorescence staining to assess the localisation of C5aR1 protein in hypoxia. Consistent with the flow cytometry results, we observed increased intracellular C5aR1 protein expression under hypoxia (Figure 4G). The increased C5aR1 protein did not appear to localise to specific intracellular organelles (Supplementary figure S4E-S4I). Furthermore, as our results suggested that C5aR1 expression was induced in a UPR-dependent manner under hypoxia, we measured intracellular C5aR1 protein levels in the presence of UPR inducers. As expected, the intracellular C5aR1 levels were increased in the presence of UPR inducers thapsigargin or tunicamycin, suggesting that ER stress specifically increases intracellular C5aR1 pools (Figure 4H and Supplementary figure S4J).

Accumulating evidence has shown that GPCRs are internalised after activation ^46,47^ and hypoxia broadly induces endocytosis in cancer cells ^48,49^. We therefore examined whether endocytosis is involved in regulating C5aR1 protein localisation in hypoxia. In the presence of endocytosis inhibitor, Dynasore, cell membrane C5aR1 levels were increased under hypoxia while not affecting the total C5aR1 levels (Figure 4I and Supplementary figure S4K). Also, there were no changes in C5aR1 expression observed upon Dynasore treatment under normoxia (Figure 4I). These results suggest that hypoxia-induced endocytosis contributes to the shift in increased C5aR1 pools from membrane to intracellular compartments. Interestingly, the depletion of C5 showed no changes in the dynamics of C5aR1 localisation under hypoxia (Figure 4J), suggesting that increased autocrine receptor activation on the membrane (via autocrine C5a ligation) is not likely to drive intracellular localisation of C5aR1 under hypoxia. Taken together, these findings suggest that UPR-induced C5aR1 is internalised and accumulates by endocytosis under hypoxia, and that hypoxia-induced intracellular C5aR1 plays a role in the cellular adaptation to ER stress under hypoxia. Importantly, intracellular C5aR1 can be used as a new therapeutic target for resistant cancer cells in the hypoxic TME (Figure 4K).

## Discussion

In this study, we show that intracellular C5aR1 expression increases in a UPR-dependent manner in various cancer cells to mediate their adaptation to hypoxic stress. Importantly, we demonstrate that genetic and pharmacological inhibition of C5aR1 decreases cancer cell survival under severe hypoxia. Upon pharmacological inhibition of C5aR1, we observe the most significant effects on cell survival following treatment with Avacopan, an approved cell permeable potent and selective small molecule inhibitor of C5aR1 ^50,51^. These findings reveal a new mechanism by which the dysregulated C5a/C5aR1 axis contributes to tumour cell survival in the hypoxic TME and provide new insights into the most effective pharmacological targeting of the C5a/C5aR1 axis. Importantly, the hypoxia-induced shift in C5aR1 localisation cautions against the use of therapeutic strategies with limited cell membrane permeability/internalisation potential.

Accumulating evidence has shown that the complement system is dysregulated in the TME ^8,9^ and this is at least in part attributed to tumour hypoxia ^52–54^. For example, a recent report indicates that hypoxia-dependent increases in C3 and C3aR expression in GBM cell lines enhance tumour growth via crosstalk with macrophages ^55^. In agreement with other studies ^25–27^, we observed that C5aR1 protein is highly expressed in tumour tissues compared with normal tissues. In addition, our *in vitro* and *in vivo* data, supported by analysis of patient data, demonstrate that increased C5aR1 expression in the hypoxic TME is mediated by the ER-stress driven UPR. These findings indicate that C5aR1 can be used as a new therapeutic target in tumours with increased hypoxia and ER stress and associated poor outcomes ^23,24^. Using UPR inhibitors and in loss-of-function experiments for transcription factors such as XBP1, ATF4 and ATF6 we find that C5aR1 expression is complementarily regulated by multiple UPR pathways; highlighting the strong dependence of tumour cells on UPR-dependent C5aR1 signalling. It remains unclear whether C5aR1 expression is directly regulated by these transcription factors, but public ChIP data indicate that some of the UPR-related transcription factors are likely to bind to the C5aR1 gene locus. Since C5aR1 is a GPCR, previous studies have shown the function of this receptor on the cell membrane ^8,10,41,42^. Here, we demonstrate that ER stress-induced C5aR1 is internalised and accumulates intracellularly in cancer cells via endocytosis. Our findings are consistent with recent reports showing that C5 can be cleaved intracellularly in multiple cell types to promote C5a/C5aR1 signalling and tumour progression ^12,43^. Of note these studies did not investigate the impact of physiologically relevant conditions of the TME (such as hypoxia) on C5a/C5aR1 signalling. Here, we demonstrate that inhibition of C5a/C5aR1 signalling either genetically or with inhibitors with high cell permeability most effectively increases autophagy and apoptosis following hypoxic stress, leading to cancer cell death. These findings support, although indirectly, the notion that intracellular C5a/C5aR1 signalling axis mediates cancer cell adaptation to hypoxic stress. Adaptation to stress requires mounting a biological response allowing the cell to return to homeostasis. We propose that ER-stress induced intracellular C5aR1, may allow the cancer cell to adapt to this stress by limiting autophagy and apoptosis, thereby promoting a return to homeostasis and cell survival. Interestingly, a recent report has shown that cell membrane C5aR1 induces autophagy and apoptosis in microglia, while these effects are reduced by inhibiting cell membrane C5aR1 with PMX205 ^56^. Together with this report, our finding that hypoxia-induced C5aR1 suppresses autophagy and apoptosis in cancer cells, supports the notion that C5aR1 plays a different role in normal and tumour tissues. These different roles might be explained by differences not only in the expression of C5aR1 between malignant and non-malignant tissue, as shown in our TMA analysis, but also in homeostatic regulatory functions based on differences in the intracellular and extracellular localisation of C5aR1 under different microenvironment conditions.

Our findings therefore provide a better understanding of the molecular mechanisms underlying dysregulated expression and function of C5aR1 under physiologically relevant TME conditions associated with poor outcome. These findings are of translational value since they can inform the choice of the most appropriate pharmacological approach to use in therapy resistant tumours with high levels of hypoxia and/or ER stress.

## Materials and Methods

### Cell lines

Adult male colorectal adenocarcinoma cell lines, HCT116 and RKO, adult female ovarian adenocarcinoma cell line SKOV3, adult male prostate adenocarcinoma cell lines, DU145 and PC3, and adult female brain glioblastoma cell line LN-229 were purchased from ATCC**^®^**. HIF1α- and p53-deficient HCT116 cells were generated as described previously ^36,57–59^. All cells were cultured in DMEM, except PC3 (cultured in RPMI) containing 10% FBS, 100 U/mL penicillin, and 100 μg/mL streptomycin. Cells were incubated in a standard humidified incubator at 37^ο^C and 5% CO_2_ for the normoxic culture, in Ruskin INVIVO_2_ 400 (Ruskinn Technology Limited) for hypoxic culture at 1% O_2_, or in BACTRON II anaerobic chamber (Shel lab) for hypoxic culture at <0.1% O_2_. For hypoxic culture at <0.1% O_2_, cells were plated on glass dishes. All cell lines were routinely tested and verified mycoplasma free using MycoAlert™ PLUS Mycoplasma Detection Kit (Lonza Bioscience, LT07-703, LT07-518).

### SiRNAs and treatments

SiRNA were transiently transfected with Lipofectamine®RNAiMax transfection reagent (Thermo Fisher Scientific, 13778) according to the manufacturer’s instructions. All siRNAs against XBP1 (L-009552-00), ATF4 (L-005125-00), ATF6 (L-009917-00), C5aR1 (L-005442-00), C5 (L-007819-00), or non-targeting RNAi negative control (Scramble, D-001810-10) were purchased from Dharmacon. C5AR1_OHu107216C_pcDNA3.1(+)-C-eGFP (pcDNA3.1/C5aR1-GFP) or empty vector were previously purchased from GenScripts and transfected with Lipofectamine 3000 (Thermo Fisher Scientific, L3000) according to the manufacturer’s instructions ^6^. CoCl_2_ (Sigma-Aldrich, 15862), thapsigargin (Thermo Fisher Scientific, T7458), tunicamycin (Sigma-Aldrich, T7765), and transferrin conjugated with Alexa Fluor 594 (Thermo Fisher Scientific, T13343) were used in this study. Before hypoxic culture, cells were pretreated with IRE1α inhibitor (4μ8c, Sigma-Aldrich, SML0949), PERK inhibitor (AMG PERK 44, Tocris, 5517), ATF6 Inhibitor (Ceapin-A7, Sigma-Aldrich, SML2330), and Dynasore (Sigma-Aldrich, D7693) for 1 hr, or with C5aR1 antagonists, PMX205 (Tocris, 5196), JPE-1375 (MedChem Express, HY-148141) and Avacopan (Cayman Chemical, CAY36639) for 8 hr, respectively.

### Spheroids

Spheroids were produced as previously described ^60^. Briefly, HCT116 spheroids were grown in Ultra Low Attachment U-bottom plates (Grenier Bio-One, #650970) to an average diameter of 500 μm, which was confirmed with a GelCount Colony Counter (Oxford Optronix). Spheroids were treated with 200 μM EF5 (Merk Millipore, # 152721-37-4) for 6 hours before fixation in 4% paraformaldehyde overnight, incubated in 30% sucrose for a minimum of 2.5 hours, and then embedded in O.C.T. Compound (VWR, TissueTek) for cryosectioning. All sectioning was conducted on a cryostat (Leica Biosystems) at -22^ο^C and spheroid sections were taken at 5 μm.

### Animal studies

Tumour-bearing mice were prepared by subcutaneously transplanting suspensions of 1.5 x 10^6^ HCT116 cells in a 1:1 ratio of 1x PBS and Matrigel (Corning, #354234) in CD-1 nude female mice at age 55–70 days. Tumour size was measured with calipers using the formula V = (**a** x **b**^2^)/2, where **a** was the largest and **b** was the smallest perpendicular diameter, respectively. Tumours ranged from 100-200 mm^3^ and were fixed in 4% paraformaldehyde for 5 hours, incubated in 30% sucrose overnight, embedded in O.C.T. Compound, and cryosectionned at 10 μm per section. Mice were euthanised using a Schedule 1 method under the UK Animals [Scientific Procedures] Act 1986 [ASPA]. Animal Research Reporting of In Vivo Experiments (ARRIVE) guidelines were used.

### Tissue microarray

Tissue microarrays were assembled by the Oxford Radcliffe Biobank (ORB)/Oxford Centre for Histopathology Research (OCHRe) team. Samples were selected from patients who had been previously recruited to ORB and/or to the Genomics England 100,000 Genomes pilot project. With the help of Oxford University Hospitals pathologists, areas of tissue in diagnostic blocks were selected to obtain punches of areas of interest. The project was covered by ORB ethical approval 19/SC/0173 and TMA blocks were placed in the ORB TMA collection for use by approved research projects.

### Immunohistochemistry

Immunohistochemistry was performed as previously described ^6^. Briefly, frozen sections of spheroids or xenografts, or formalin-fixed paraffin-embedded sections (after dewaxing and hydration) of tissue microarray were subjected to antigen retrieval and blocking with 0.05% Tween 20 Citrate Buffer (Sigma-Aldrich, C9999) and Serum-Free Protein Block (DAKO, X0909) according to the manufacturer’s instructions, respectively. Sections were then stained using the following primary antibody; anti-C5aR1 rabbit polyclonal antibody (Abcam, catalogue ab59390), anti-C5aR1 mouse monoclonal antibody (BD Biosciences, 550493), anti-BIP rabbit monoclonal antibody (Cell Signaling, 3177) and anti-EF5 mouse monoclonal antibody conjugated with Alexa Fluor 488, (EF5 Hypoxia Detection Kit, Sigma-Aldrich, EF5-30A4). As second antibodies, Alexa Fluor 488 or 594 goat anti-rabbit IgG and Alexa Fluor 594 donkey anti-mouse IgG (Thermo Fisher Scientific) for the immunohistofluorescence staining were used. EnVision+ System (Dako) was used for DAB staining. As counterstains, DAPI and hematoxylin were used for the immunohistofluorescence and DAB staining, respectively. For immunohistofluorescence staining, the section was scanned and analysed using The Nikon NiE microscope and ImageJ software. For DAB staining, the whole section was scanned and analysed using the Aperio CS scanner (Aperio Technologies) and QuPath software.

### Colorimetric cell viability assay

Cells were subjected to the cell viability assay using MTT Reagent (Sigma-Aldrich, M2128). The cell viabilities post-hypoxic or post-normoxic culture were divided by those of pre-hypoxic culture to evaluate the difference in cell viabilities between normoxic and hypoxic conditions.

### Clonogenic survival assay

The indicated cells were cultured under normoxia or hypoxia for the indicated periods following siRNA transfection and cultured for 7-10 additional days in a standard humidified incubator at 37^ο^C and 5% CO_2_. Surviving colonies were stained with Crystal Violet solution (Sigma-Aldrich, C0775). Surviving fractions were calculated, as previously described ^6,61^.

### Immunocytochemicstry

Cells were fixed with 4% PFA in each oxygen condition, and then stained as previously described ^6,62^. The following primary antibodies were used; γ-H2AX (Ser139) (Sigma-Aldrich, 05-636), RPA32/RPA2 (Cell Signaling, 2208), CD88 (BD Biosciences, 550493), LAMP1 (Cell Signaling, 9091), AIF (Cell Signaling; 5318), PDI (Cell Signaling, 3501), EEA1 (Cell Signaling, 3288), RCAS1 (Cell Signaling, 12290). As secondary antibodies, Alexa Fluor 594 goat anti-rabbit IgG and Alexa Fluor 488 or 594 donkey anti-mouse IgG (Thermo Fisher Scientific) were used. Alexa Fluor 647 Phalloidin (Thermo Fisher Scientific, A22287) and DAPI stain (Thermo Fisher Scientific, P36962) were added to each coverslip if necessary, and imaged on the Zeiss LSM 710 Confocal Microscope.

Apoptosis assessment by morphology was carried out as previously described ^6,11,59^. Briefly, cells with fragmented DNA were recognised as apoptotic cells and the percentage of the apoptosis was calculated by the ratio of the apoptotic cells to the total cells per field (with typically at least 100 cells counted for each group).

### Immunoblotting

Cells were harvested in UTB (9 M urea, 75 mM Tris-HCl pH 7.5 and 0.15 M β-mercaptoethanol), and then sonicated. The cell lysate was subjected to immunoblotting using the following primary antibodies; β-actin (Sigma-Aldrich, A5441), HIF-1α (BD Biosciences, 610959), BiP (Cell Signaling, 3177), p62 (Cell Signaling, 88588), LC3B (Cell Signaling, 3868). LI-COR Odyssey imaging system was used. Experiments were repeated at least three times and representative blots were shown unless otherwise noted.

### Flow cytometry analyses

Cells were fixed with 4% PFA. Samples were either permeabilised (with methanol for 30 min on ice) for total C5aR1 or not permeabilised for cell membrane C5aR1 expression analysis. Cells were blocked in 2% BSA PBS, incubated for 1 hr on ice with BD OptiBuild BV421 Mouse Anti-Human CD88 antibody (BD Biosciences, 742315) or BD Horizon BV421 Mouse IgG1, k Isotype Control (BD Biosciences, 562438), and then subjected to flow cytometry.

For cell cycle analysis, samples were incubated in 5.6 μg/mL PureLink RNase A (Thermo Fisher Scientific, 12091) and 89 μg/mL propidium iodide solution (Thermo Fisher Scientific, P3566) in PBS for 10 min at room temperature, and subjected to cell cycle analysis as previously described ^6^.

Data acquisition was carried out using CytoFLEX (Beckman Coulter Inc.). Data analysis was carried out using FlowJo Software (BD Biosciences).

### qRT-PCR

Total RNA was extracted using TRI Reagent (Sigma-Aldrich) and subjected to reverse transcription using Verso cDNA Synthesis Kit (Thermo Scientific). Quantitative real-time PCR was performed using SYBR Green PCR Master Mix kit (Applied Biosystems) in StepOnePlus™ Real-Time PCR system (Applied Biosystems) to quantify mRNA levels of the indicated genes. Relative mRNA levels were normalised to human ACTB or 18S as an internal control and calculated using a 2^-ΔΔCt^ method. All primer sequences are listed in Supplemental Table 1.

### TCGA analysis

Publicly available patient RNA-seq datasets were obtained from The Cancer Genome Atlas (TCGA) via the GDC Data Portal (https://portal.gdc.cancer.gov). RNA-seq datasets for colorectal (TCGA-COAD and TCGA-READ), glioblastoma (TCGA-GBM), prostate (TCGA-PRAD) and ovarian (TCGA-OV) were downloaded using the *TCGAbiolinks* (ver. 2.30.0) package and analysed in R. Unstranded raw gene expression counts for each cohort along with gene and patient metadata was extracted from all downloaded datasets for analysis. Unstranded raw expression counts for all datasets were transformed using the variance stabilising transformation (VST) method implemented with the *DESeq2* function *vst*, subsequently subsetting the expression matrix for tumour only expression data using patient metadata.

Correlation analysis amongst C5aR1 and other identified gene signatures for hypoxia and UPR was performed using the transformed gene expression data for each cohort. Pearson’s correlation coefficient was used to assess correlation strength and directionality of relationship amongst C5aR1 and gene signatures in all datasets. All genes for identified gene signatures were listed in Supplemental Table 2^63–66^.

Kaplan-Meier curves for overall survival and disease-free survival of TCGA patients with the indicated cancers were obtained from Gene Expression Profiling Interactive Analysis 2 (GEPIA2) (http://gepia2.cancer-pku.cn). Samples were categorised by the expression levels of C5aR1 as either “high” or “low” group. *P* values were calculated by log-rank test.

### Statistics

The experimental data were shown using individual biological replicates (and with the technical replicates if available) according to the literature ^67^. The data obtained from the same experiment is shown in the same coloured and shaped plots; circle for the fist, triangle for the second, square for the third, rhombus for the fourth experiment. Statistical analysis was carried out using GraphPad Prism software (version 10). *p* values were used to determine significance of differences and calculated using biological replicates, and *p* values less than 0.05 were considered significant. Two-tailed paired or unpaired Student’s t test and one- or two-way ANOVA were used as appropriate and as described in each figure legend.

### Study approval

All animal experiments were conducted under a project licence (PPL30/2922), which was approved by the Oxford University Animal Welfare and Ethical Review Body (AWERB) and issued by the UK Home Office under the UK Animals (Scientific Procedures) Act of 1986.

Use of the tissue microarray was approved by the National Research Ethics Service in the UK (REC ref. 15/EE/0241, IRAS ref 169363, project 23/A007). Written-informed consent was obtained from all patients.

## Data availability

All data can be accessed from the corresponding author upon request.

## Author contributions

TS., KL., IC., DJM., HC. and MMO provided methodology. TS and MMO wrote the original draft of the manuscript. TS, KL., IC., DJM., HC., NM, GRB, GSH, EMH and MMO reviewed and edited the manuscript. TS and MMO conceptualised the study. TS, KL., IC., DJM., HC., and MMO provided investigation and formal analysis. NM, GRB, GSH, EMH and MMO provided resources. GSH, EMH and MMO provided supervision. TS, GSH, EMH and MMO acquired funding.

## Acknowledgements

This work was supported by MRC (MC_UU_00001/10), NIH Grant CA257907-01A1, the Academy of Medical Sciences (AMS), the Wellcome Trust, the Government Department of Business, Energy and Industrial Strategy (BEIS), the British Heart Foundation and Diabetes UK through an AMS Springboard (REF:SBF008/1156, awarded to MMO) and Cancer Research UK (CRUK) grant number C5255/A18085, through the Cancer Research UK Oxford Centre. This work was also supported by CRUK grant number CTRQQR-2021\100002 through the Oxford Centre for Early Cancer Detection and Cancer Research UK Oxford Centre. TS is supported by Japan Society for the Promotion of Science (JSPS) Overseas Research Fellowships, Takeda Science Foundation Fellowship, Kanzawa Medical Research Foundation Fellowship and the British Council Japan Association (BCJA) Scholarship Scheme.

We acknowledge the Oxford Centre for Histopathology Research and the Oxford Radcliffe Biobank, which are supported by the University of Oxford, the Oxford CRUK Cancer Centre and the NIHR Oxford Biomedical Research Centre (Molecular Diagnostics Theme/Multimodal Pathology Subtheme), and the NIHR CRN Thames Valley network. We would also like to acknowledge the Tissue Histopathology Laboratory (University of Oxford) and, in particular, Leticia Campo Urriza and Molly Browne.

