## Supplementary Information for "UPR-induced intracellular C5aR1 promotes adaptation to the hypoxic tumour microenvironment by regulating tumour cell fate"

**Supplementary Figure S1**

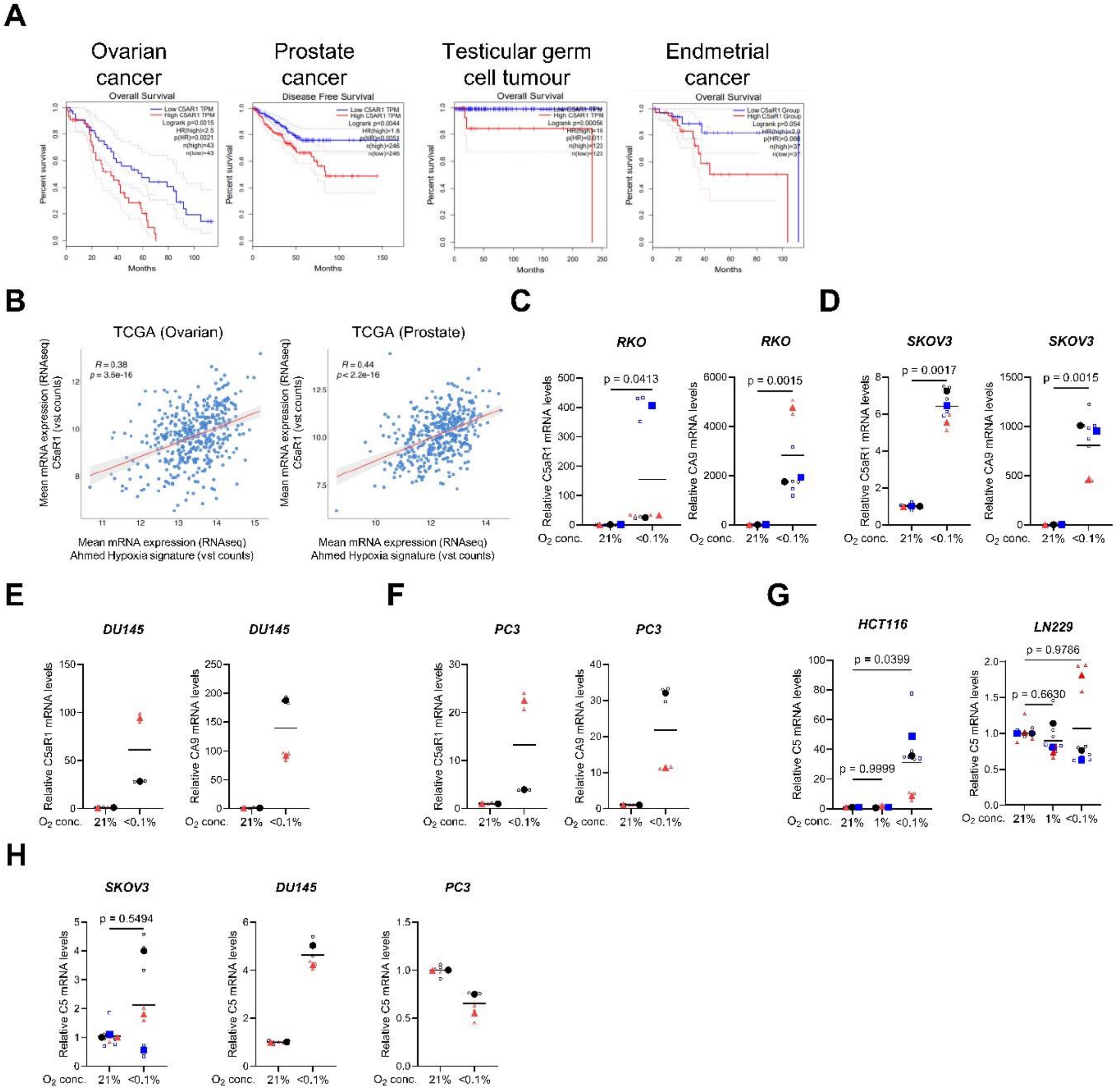

**Supplementary Figure S1. C5aR1 is highly expressed in hypoxic regions and is associated with poor outcome in various cancers**

**(A)** GEPIA Kaplan-Meier curve for overall survival of TCGA ovarian and endometrial cancer and testicular germ cell tumour patients and disease free survival of TCGA prostate cancer patients with high (red) or low (blue) C5aR1 mRNA expression levels is shown. (http://gepia.cancer-pku.cn).

**(B)** Pearson's correlation between C5AR1 mRNA expression and Ahmed Hypoxia signature in TCGA ovarian (left) and prostate (right) cancer samples. *R* score and *p* value are shown.

**(C and D)** RKO (C) and SKOV3 (D) cells were cultured under normoxia or hypoxia (<0.1% $O_2$) for 24 hours (hr) and subjected to qRT-PCR. n=3.

**(E and F)** DU145 (E) and PC3 (F) cells were cultured under normoxia or hypoxia (<0.1% $O_2$) for 24 hr, and subjected to qRT-PCR. n=2.

**(G and H)** C5 mRNA levels in the experiments of Figure 1D and 1E (G), and Supplementary Figure S1D-S1F (H) were shown. n=3 for HCT116, LN229 and SKOV3 cells, and n=2 for DU145 and PC3 cells.

Individual biological replicates (large points) represent the average of the technical replicates (small points). *p* values were calculated using biological replicates by one-way two-tailed paired Student's t test (C, D, and H) and ANOVA with Dunnett's test (G).

#### Supplementary Figure S2

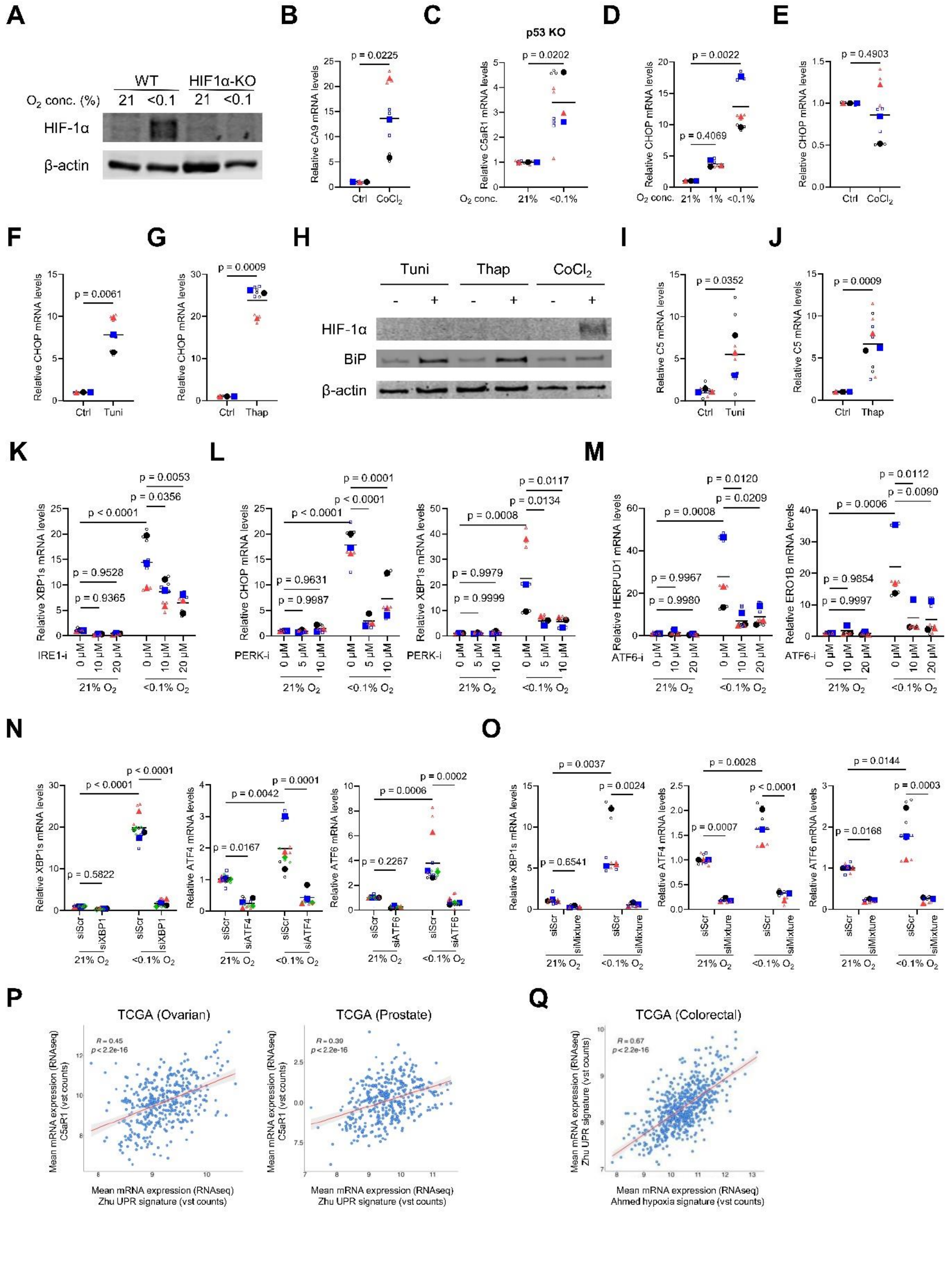

**Supplementary Figure S2. ER stress induces C5aR1 expression in cancer cells under hypoxia (<0.1% $O_2$)**

For the whole figure: Individual biological replicates (large points) represent the average of the technical replicates (small points). *p* values were calculated using biological replicates by two-tailed paired Student's t test (B, C and E-G), two-tailed unpaired Student's t test (I and J), two-way ANOVA with uncorrected Fisher's LSD test (K-O), or one-way ANOVA with Dunnett's test (D).

**(A)** HIF-1α- KO and WT HCT116 cells were cultured under normoxia or hypoxia (<0.1% $O_2$) for 24 hr, and subjected to Immunoblotting.
**(B)** CA9 mRNA levels in the experiments of Figure 2B were shown. n=3.
**(C)** p53- KO HCT116 cells were cultured under normoxia or hypoxia (<0.1% $O_2$) for 24 hr, and subjected to qRT-PCR. n=3.
**(D)** HCT116 cells were cultured in the indicated conditions for 24 hr, and subjected to qRT-PCR. n=3.
**(E-J)** HCT116 cells were cultured in the same conditions as Figure 2B-D, and subjected to qRT-PCR (E-G, I and J) and Immunoblotting (H). n=3.
**(K)** XBP1s (downstream gene of IRE1) mRNA levels in the experiments of Figure 2E were shown. n=3.
**(L)** CHOP and XBP1s (downstream genes of PERK) mRNA levels in the experiments of Figure 2F were shown. n=3.
**(M)** HERPUD1 and ERO1B (downstream genes of ATF6) mRNA levels in the experiments of Figure 2G were shown. n=3.
**(N)** XBP1s, ATF4 and ATF6 mRNA levels in the experiments of Figure 2I are evaluated to analyse the knockdown efficiencies. n=4.
**(O)** XBP1s, ATF4 and ATF6 mRNA levels in the experiments of Figure 2J are evaluated to analyse the knockdown efficiencies. n=3
**(P)** Pearson's correlation of C5AR1 mRNA expression and Xhu UPR signature in TCGA ovarian (left) and prostate (right) cancer samples. *R* score and *p* value are shown.
**(Q)** Pearson's correlation between Xhu UPR and Ahmed Hypoxia signature in TCGA colorectal cancer samples. *R* score and *p* value are shown.

#### Supplementary Figure S3

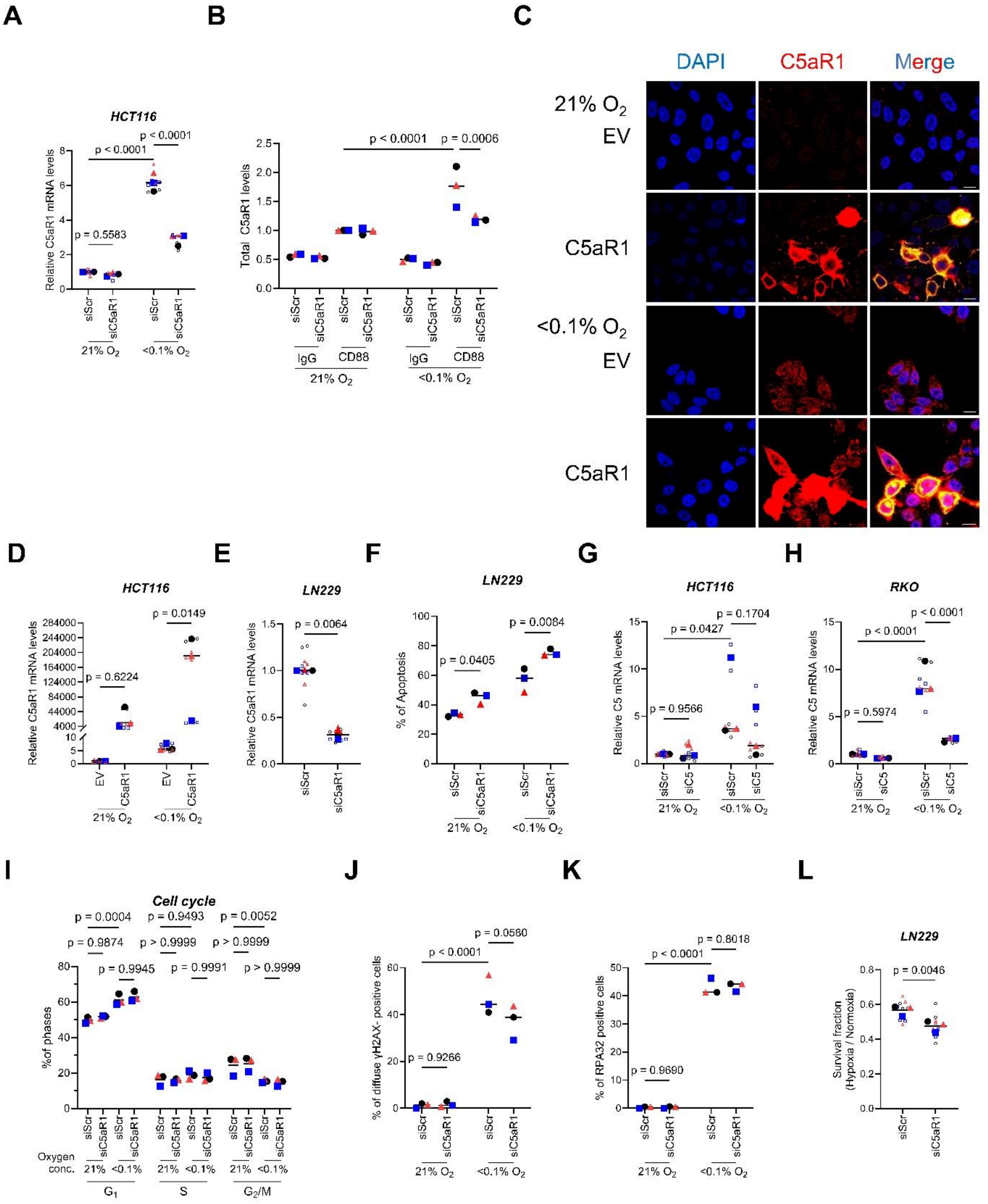

**Supplementary Figure S3. Hypoxia-induced C5aR1 mediates cellular adaptation to hypoxic stress by regulating cancer cell death**
For the whole figure: Individual biological replicates (large points) represent the average of the technical replicates (small points). *p* values were calculated using biological replicates (large points) by two-way ANOVA with uncorrected Fisher's LSD test (A, B, D, F, G, H, J and K), two-tailed paired Student's t test (E and L), or two-way ANOVA with Sidak's multiple comparison test (I).

**(A and B)** HCT116 cells were transfected with either siRNA against C5aR1 (siC5aR1) or scramble siRNA (siScr) for negative control, cultured under normoxia or hypoxia (<0.1% $O_2$) for 24 hr, and subjected to qRT-PCR (A) and Flow Cytometry following permeabilisation (B). n=3.
**(C)** RKO cells were transfected with either pcDNA3.1/C5aR1-GFP (C5aR1) or its empty vector (EV), cultured under normoxia or hypoxia (<0.1% $O_2$) for 24 hr, and subjected to Immunocytochemistry. C5aR1 (red) or DAPI (blue). Scale bar, 10 µm.
**(D)** HCT116 cells were transfected with either pcDNA3.1/C5aR1 (C5aR1) or its empty vector (EV), cultured under normoxia or hypoxia (<0.1% $O_2$) for 24 hr, and subjected to qRT-PCR. n=3.
**(E)** LN229 cells were transfected with either siC5aR1 or siScr, cultured under normoxia for 72 hr and subjected to qRT-PCR. n=3.
**(F)** LN229 cells were transfected with either siC5aR1 or siScr, cultured under normoxia or hypoxia (<0.1% $O_2$) for 16 hr and subjected to apoptosis assay. n=3.
**(G and H)** HCT116 (G) and RKO (H) cells were transfected with siRNA against C5 (siC5) or siScr, cultured under normoxia or hypoxia (<0.1% $O_2$) for 24 hr, and subjected to qRT-PCR. n=3.
**(I)** HCT116 cells were transfected with either siC5aR1 or siScr, cultured under normoxia or hypoxia (<0.1% $O_2$) for 16 hr, and subjected to Flow Cytometry for cell cycle analysis. n=3.
**(J and K)** After transfection with either siC5aR1 or siScr, HCT116 cells were subjected to Immunocytochemistry for γH2AX and RPA32 focus assay following hypoxic treatment for 24 and 8 hr, respectively. n=3.
**(L)** LN229 cells were transfected with either siC5aR1 or siScr, cultured under normoxia or hypoxia (<0.1% $O_2$) for 16 hr and subjected to clonogenic survival assay. n=3.

### Supplementary Figure S4

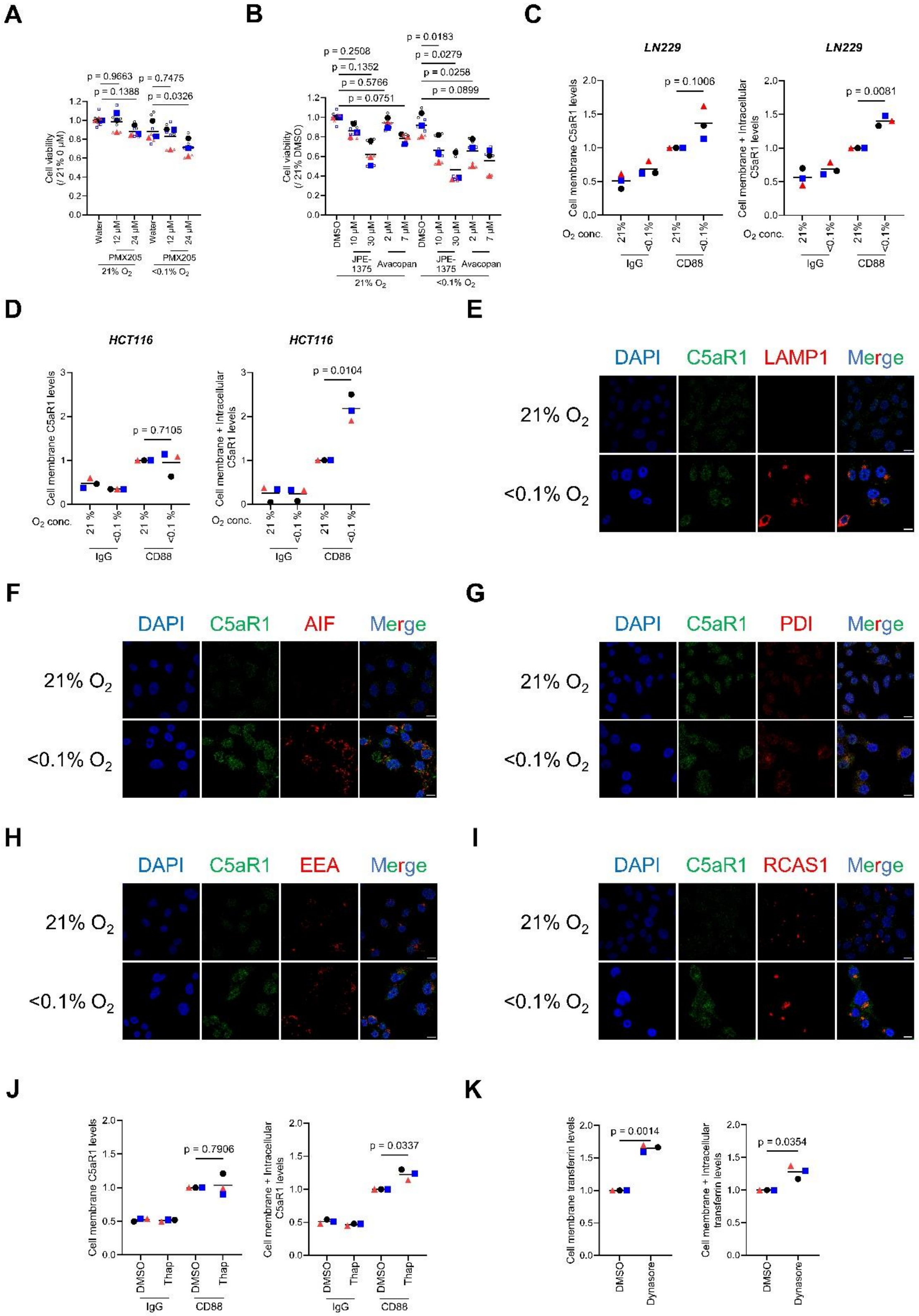

**Supplementary Figure S4. Pharmacologically targeting intracellular C5aR1 results in reduced tumour cell viability with enhanced autophagy and apoptosis**
For the whole figure: Individual biological replicates (large points) represent the average of the technical replicates (small points). *p* values were calculated using biological replicates (large points) by one-way ANOVA with Dunnett's test (A and B), two-way ANOVA with uncorrected Fisher's LSD test (C and D), and two-tailed paired Student's t test (J and K).

**(A and B)** HCT116 cells were pretreated for 8 hr with the indicated dose of C5aR1 abtagonists, PMX205 (A), JPE-1375 and Avacopan (B), cultured under normoxia or hypoxia (<0.1% $O_2$) for 24 hr, then cultured under normoxia for 40 hr, and subjected to cell viability assays. n=3.
**(C and D)** LN229 (C) and HCT116 (D) cells were cultured under normoxia or hypoxia (<0.1% $O_2$) for 16 hr and 24 hr, respectively, and subjected to FACS, with (right) or without (left) permeabilisation. n=3.
**(E-I)** RKO cells were cultured under normoxia or hypoxia (<0.1% $O_2$) for 24 hr, and subjected to triple immunofluorescence. C5aR1 (green), DAPI (blue), or Organelle makers (red); LAMP for lysosome (E), AIF for mitochondria (F), PDI for endoplasmic reticulum (G), EEA for endosomal membrane (H), RCAS1 for Golgi (I). Scale bar, 10 µm.
**(J)** HCT116 cells were treated with 2 µM thapsigargin (Thap) or its vehicle (DMSO) for 16 hr and then subjected to FACS with (right) or without (left) permeabilisation. n=3.
**(K)** HCT116 cells were cultured under normoxia in the presence of 100 µM Dynasore and 25 µg/ml transferrin conjugated with Alexa Fluor 594, and then subjected to FACS with (right) or without (left) permeabilisation. n=3.

**Supplemental Table 1.** List of primers used in qRT-PCR experiments.

| target gene | forward primer (5'-3') | reverse primer (5'-3') |
|---|---|---|
| ACTB | ACATCCGCAAAGACCTCTACG | TTGCTGATCCACATCTGCTGG |
| 18S | GTGGAGCGATTTGTCTGGTT | ACGCTGAGCCAGTCAGTGTA |
| C5aR1 | TCCTTCAATTATACCACCCCTGA | ACGCAGCGTGTTAGAAGTTTTAT |
| CA9 | CTTGGAAGAAATCGCTGAGG | TGGAAGTAGCGGCTGAAGTC |
| C5 | CTCCTCAGGCCATGTTCATT | TCTTTTGGCTGGCTTCAAGT |
| CHOP | GGGAGCTGGAAGCCTGGTA | CCCCCATTTTCATCTGAAGACA |
| XBP1s | TGCTGAGTCCGCAGCAGGTG | GCTGGCAGGCTCTGGGGAAG |
| HERPUD1 | AACGGCATGTTTTGCATCTG | GGGGAAGAAAGGTTCCGAAG |
| ERO1B | AATCTGAAGCGACCTTGTCC | GCCCAGCTTTTATTCCAACC |
| ATF4 | GGAGATAGGAAGCCAGACTACA | GGCTCATACAGATGCCACTATC |
| ATF6 | CAGACAGTACCAACGCTTATGCC | GCAGAACTCCAGGTGCTTGAAG |

**Supplemental Table 2.** List of genes for gene signatures used in TCGA analysis

| ***Ahmed hypoxia signature*** |
| --- |
| ANXA1 |
| CALD1 |
| CP |
| IGFBP2 |
| IGFBP5 |
| LOX |
